# Evolutionary Action Score of *TP53* Mutations: Integrated Clinico-pathologic And Protein Structural Analysis in Myelodysplastic Syndromes

**DOI:** 10.1101/2020.07.08.194365

**Authors:** Rashmi Kanagal-Shamanna, Guillermo Montalban-Bravo, Panagiotis Katsonis, Koji Sasaki, Caleb A. Class, Christopher Benton, Elias Jabbour, Kelly S. Chien, Rajyalakshmi Luthra, Carlos E. Bueso-Ramos, Tapan Kadia, Michael Andreeff, Nicholas Short, Naval Daver, Mark J. Routbort, Joseph D. Khoury, Keyur Patel, Irene Ganan-Gomez, Yue Wei, Gautam Borthakur, Farhad Ravandi, Kim-Anh Do, Kelly A. Soltysiak, Olivier Lichtarge, L. Jeffrey Medeiros, Hagop Kantarjian, Guillermo Garcia-Manero

## Abstract

To determine the impact of *TP53* mutations on the phenotype and outcome of myelodysplastic syndromes, we quantified the deleterious effects of missense *TP53* mutations using the computationally-derived evolutionary action score (higher score indicates worse impact), based on the phylogenetic divergence of the sequence position and amino acid change perturbation, and correlated with clinico-pathologic-genomic features in 270 newly-diagnosed *TP53*-mutant patients primarily treated with hypomethylating agents. Using recursive partitioning and regression trees, we identified a subset of patients with low-EAp53 mutations (≤52) with improved overall survival (OS) (n=17, 6%) compared to high-EAp53 (n=253, 94%) [median OS, 48 vs. 10 months; p=0.01]. Compared to high-EAp53, low-EAp53 patients had lower cytogenetic complexity, lower TP53 protein expression, lacked multi-allelic *TP53* alterations, but had more somatic mutations in other genes. There was no difference in median *TP53* variant allele frequency or distribution of R-IPSS. 3D-protein modeling showed clustering of poor-outcome mutations, indicating structural location influences outcome.

## INTRODUCTION

*TP53* mutations are present in about 10% of MDS patients and are associated with complex karyotype, deletion (5q) and an adverse prognosis^1, 2, 3, 4, 5, 6, 7, 8, 9^. Rarely, *TP53* mutations can be present in low-risk MDS^10, 11^. Although patients with *TP53*-mutated MDS generally have a poor outcome, prognosis can be heterogeneous depending on mutation characteristics such as variant allele frequency (VAF) and associated parameters, such as the presence of complex karyotype (CK) and concurrent 17p deletion^5, 7, 9, 10, 11, 12, 13, 14, 15^. We and others have shown that complex karyotype defines worse survival in patients with *TP53* mutated MDS, as a consequence of the genomic instability acquired by the mutant clone. Accordingly, *TP53* mutated MDS patients without a complex karyotype have significantly better outcomes, especially in the setting of low VAF ^16^. In addition, a recent study by Bernard, *et al* demonstrated that mono-allelic *TP53* mutations were associated with significantly better outcomes compared to multi-allelic *TP53* abnormalities^17^. However, the relationship between genomic/phenotypic differences and the type of *TP53* mutations is not known.

*TP53* mutations are distributed across the entire coding region, particularly in the DNA binding domain, with less than a third of the mutations occurring in focal hotspots^18, 19, 20, 21^. *In vitro* and *in silico* studies suggest that different types of mutations lead to distinct functional consequences, with some*TP53* mutations leading to loss-of-function of the wild-type protein, exerting a dominant negative effect on the wild-type allele, and others resulting in oncogenic gain-of-function leading to cellular transformation and resistance to chemotherapy^20, 22, 23, 24, 25, 26, 27^. Hence, it is reasonable to hypothesize that the functional effects of these mutations influences disease biology and outcome, either independently or by influencing known variables such as VAF and karyotype. To date, the relationship between genomic/phenotypic differences and the type of *TP53* mutations has not been elucidated. This could be potentially important to assess the efficacy of novel therapeutic approaches that restore TP53 function^28, 29^.

The Evolutionary Action score is a computationally derived score to quantify the deleterious impact of different types of missense *TP53* mutations (EAp53). EAp53 calculates the deleterious effect of missense coding mutations based on 2 components: (1) the degree of phylogenetic divergence of the sequence position affected by the mutation, measured by the “Evolutionary Trace” approach^30, 31^, and (2) the perturbation size of the amino acid substitution of the point mutation, measured in terms of log-odds^32, 33^. The EAp53 scoring system ranges between 0 to 100, with a higher score indicating a higher impact on downstream transcription and protein function. EAp53 scoring system has been shown to be a reliable prognostic marker in patients with head and neck and colorectal cancers^25, 26, 34^ and in objective assessments^35, 36^.

In this study, we used the EAp53 scoring system to evaluate the impact of different types of missense *TP53* mutations on the phenotype, prognosis and outcome in a well-characterized cohort of 270 patients with MDS and oligoblastic acute myeloid leukemia (AML with 20-29% blasts or MDS with excess blasts in transformation). A higher EAp53 score was associated with worse outcome, and a cut-off score of 52 predicted for worse overall survival (OS). Low-EAp53 (<52) and high-EAp53 (≥52) MDS showed significant differences in clinical, genomic and cytogenetic characteristics and TP53 protein expression. 3D structural mapping of high-EAp53 mutations on TP53 protein suggested that structural location of mutations further influenced the outcome. We conclude that EAp53 score identifies prognostic subsets within *TP53*-mutant MDS patients and can facilitate a personalized therapeutic approach.

## RESULTS

### Baseline characteristics of the study group

A total of 270 MDS patients with at least one missense *TP53* mutation were identified. Patient characteristics are shown in **Table 1**. The median age was 68 years (range, 18-90). The majority of patients belonged to “very poor” revised International Prognostic Scoring System (IPSS-R) (162, 60%) category, followed by poor (61, 23%), intermediate (19, 7%), good (19, 7%) and very good (9, 3%). One hundred twenty-two (45%) were therapy-related. 219 (81%) patients had a complex karyotype. WHO diagnosis included: therapy-related myeloid neoplasm (t-MN; n=122, 46%); MDS with excess blasts (MDS-EB; 66, 24%); MDS with single lineage dysplasia (MDS-SLD) [1, 0.4%]; MDS-SLD with ring sideroblasts (MDS-SLD-RS) [2, 0.7%]; MDS with multilineage dysplasia (MDS-MLD) [22, 8%], MDS-MLD-RS [7, 2.6%]; MDS with isolated del(5q) [2, 0.7%], MDS unclassified [2, 0.7%] and AML with MDS-related changes (AML-MRC)/MDS with excess blasts in transformation (MDS-EB-T) [45, 16.7%]. Over a median follow-up of 8.1 months, the median OS was 10.3 months. Fifty-four (20%) MDS patients transformed to AML.

**Table 1.** Patient characteristics of *TP53* mutated MDS patients.

| **Characteristics** | **MDS (n=270)** |
| --- | --- |
| Age (median/ range) | 68.3 (18-90) |
| Gender: Female/ male | 102 (37%)/ 168 (62%) |
| Hemoglobin (g/dL) | 9.0 (6.4-16.2) |
| Platelet (**x10<sup>9</sup>/L**) | 49 (2-4816) |
| Absolute Neutrophil Count (**x10<sup>9</sup>/L**) | 1.1 (0-19.8) |
| PB blast% (median/ range) | 1 (0-25) |
| BM Blast% (median/ range) | 7 (0 to 28) |
| **IPSS-R categories** |  |
| 1/ 2/ 3/ 4/ 5 | 9 (3.3%)/ 19 (7%)/ 19 (7%)/ 61 (22.6%)/ 162 (60%) |
| **WHO classification** |  |
| Therapy-related myeloid neoplasm | 122 (46%) |
| MDS-SLD-RS/ MDS-MLD-RS | 2 (0.7%)/ 7 (2.6%) |
| MDS-SLD / MDS-MLD | 1 (0.4%)/ 22 (8%) |
| MDS with excess blasts (MDS-EB) | 66 (24%) |
| MDS-Unclassifiable | 3 (1.1%) |
| MDS with isolated del5q | 2 (0.7%) |
| AML-MRC (MDS-EB-T) | 45 (16.7%) |
| **Karyotype** |  |
| Diploid* | 23 (8.5%) |
| Complex* | 219 (81.1%) |
| del(5q)/ monosomy 5 | 118 (43.7%) |
| del(7q)/ monosomy 7 | 94 (34.8%) |
| del(17p)* | 93 (34.4%) |
| Monosomal karyotype | 232 (86%) |
| **Number of *TP53* mutations\*** |  |
| 1/ 2/ 3/ 4 | 189 (70%)/ 75 (28%)/ 5 (1.9%)/ 1 (0.4%) |
| ***TP53* allelic state: Monoallelic/ multiallelic** | 105 (39%)/ 165 (61%) |
| **Median VAF for *TP53* mutation** | 33.9 (1-94.4) |
| **EAp53 score** | 78.5 (4.2 – 97.94)) |
| High EA/ Low EA | 253 (94%)/ 17 (6%) |
| **Treatment regimens** |  |
| Chemotherapy-based | 17 (6.3%) |
| Chemo + HMA-based | 8 (3%) |
| Chemo + other (non-HMA-based) | 1 (0.4%) |
| HMA alone | 111 (41.1%) |
| HMA + other | 48 (17.8%) |
| Others | 3 (1.1%) |
| None | 22 (8.1%) |
| Unknown | 51 (19%) |
| Transplant | 36 (13.3%) |
| Number died | 192 (71%) |

MDS, Myelodysplastic Syndrome; VAF, Variant Allele Frequency; IPSS, R-IPSS, revised International Prognostic Scoring System; HMA, hypomethylating agent

### TP53 mutation characteristics

Only patients who had at least 1 missense *TP53* mutation were included. At the time of diagnosis, 189 (70%) patients had a single *TP53* mutation whereas 81 (30%) patients had ≥2 *TP53* mutations: 75 (28%) patients had 2, 5 (2%) patients had 3 and 1 (0.4%) patient had 4 mutations. Among the patients with ≥2 *TP53* mutations, 22 (8%) had a concurrent non-missense *TP53* mutation. Most (265, 98%) mutations were located within the DNA-binding domain (**Figure 1A**); majority (251, 93%) had mutations involving codons between 120 and 283. Concurrent del(17p) was observed in 93 (34.4%) of patients. The allelic state of *TP53* was determined based on the presence of concurrent *TP53* alterations (deletion and/or additional *TP53* mutations) as described elsewhere ^16^. A majority (165, 61%) had multi-allelic alterations of the *TP53* gene. The median VAF for *TP53* mutation was 33.9 (1-94.4). In 31 (11.5%) patients, the *TP53* VAF was <20%.

**Figure 1.**
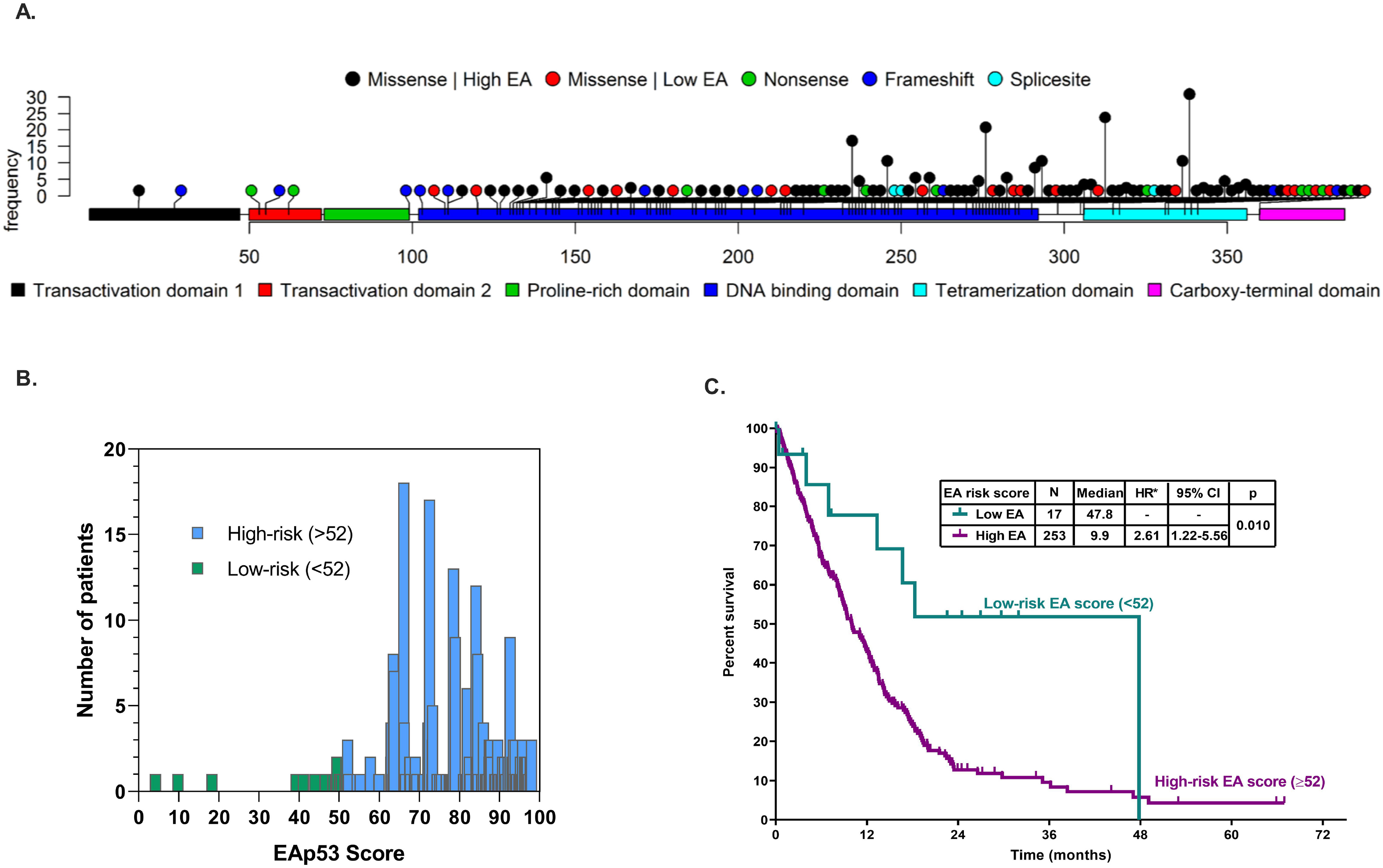
**(A)** Lollipop plot showing the frequency distribution of missense *TP53* mutations and associated concurrent non-missense mutations. **(B)** Spectrum of EAp53 scores within our MDS cohort: majority had a high (>52) EAp53 score. **(C)** By AIC (Akaike information criterion), an EAp53 score of 52 provided an optimal cut-off based on overall survival in MDS patients.

### EAp53 score and outcomes

EAp53 is a computational score to measure the deleterious impact of missense *TP53* mutations, ranging from 0-100, with higher scores indicating a higher impact on TP53 transcriptional activity and protein function and 0 indicating wild-type TP53 function. In this MDS cohort, the EAp53 scores ranged between 4.2 and 97.9, with a median of 79 (**Figure 1B**). A higher EAp53 score (as a continuous variable) correlated with worse OS (p-value: p=0.087; hazard ratio 1.06 per 10-point increase [95% CI: 1.01-1.13]). Using recursive partitioning and regression trees (RPART, based on the CART algorithm), we identified an EAp53 score >52 predicted for worse OS (**Figure 1C**). Based on this, we divided our cohort into 2 groups: low-risk EAp53 patients [defined as EAp53 ≤52; n=17 (6%)] and high-risk EAp53 patients [EAp53 >52; n=253, 94%]. The median OS for low-risk EAp53 MDS patients was 47.8 months compared to 10 months for high-risk EAp53 patients (p=0.01; HR: 2.6 [1.22-5.56]). The subset of low-risk EAp53 patients with significantly better OS, although a small number, was of great interest. We studied the association with clinico-pathologic, cytogenetic and molecular features of this subset in comparison to the high-risk patients (results presented in the subsequent section). An EAp53 score cut-off of 75, previously described in *TP53* mutated head and neck squamous cell carcinoma to show significant difference in survival, time to metastasis and response to platinum based therapies), did not show significant survival differences in MDS.

By univariate analysis, the following parameters were associated with worse OS: higher *TP53* VAF (as a continuous variable), higher number of *TP53* mutations, high-risk EAp53 (>52) group, higher IPSS-R score, presence of CK/ monosomal karyotype (MK), higher serum LDH and creatinine levels, lower platelet count, lower hemoglobin, and lower serum albumin. *TP53* allele state (mono vs. multi-allelic), del(17p), and peripheral blood (PB) blast percentage did not associate with OS. The EAp53 risk group did not affect TFS, RFS, ORR, or CR rates (**Supplemental Table 1**). By multivariable analysis, the EAp53 risk group retained the independent predictive value for OS along with IPSS-R score, serum albumin, and serum bilirubin. Neither *TP53* VAF nor the number of *TP53* mutations was independently prognostic. We excluded the presence of a complex karyotype in the model due to a strong association with EAp53 score (**Table 2**). EAp53 risk category was the only independent predictor of AML transformation.

**Table 2.** Multivariable model incorporating clinical and mutational characteristics of *TP53* mutation OS.

| Parameters | HR | lower .95 | upper .95 | p-value |
| --- | --- | --- | --- | --- |
| Number of TP53 mutations | 0.8091 | 0.5207 | 1.2571 | 0.34604 |
| TP53 VAF | 1.0028 | 0.9936 | 1.012 | 0.55441 |
| **EAp53 (low vs. high)** | 5.0643 | 1.4928 | 17.1801 | **0.00925** |
| **IPSS-R score** | 1.2524 | 1.1184 | 1.4025 | **9.74E-05** |
| Serum LDH | 1 | 0.9996 | 1.0004 | 0.87182 |
| **Serum Albumin** | 0.57 | 0.362 | 0.8978 | **0.0153** |
| Fibrinogen | 1.0015 | 0.9999 | 1.003 | 0.06037 |
| Serum creatinine | 1.0547 | 0.7602 | 1.4634 | 0.74993 |
| **Serum Bilirubin** | 1.6136 | 1.0381 | 2.5082 | **0.03351** |

**AML transformation**
| Parameters | HR | lower .95 | upper .95 | p-value |
| --- | --- | --- | --- | --- |
| TP53 VAF | 0.999 | 0.9871 | 1.011 | 0.8668 |
| **EAp53 (low vs. high)** | 7.4143 | 0.931 | 59.049 | **0.0584** |
| IPSS-R score | 1.1099 | 0.8702 | 1.416 | 0.401 |
| Complex cytogenetics | 3.0833 | 0.6095 | 15.596 | 0.1734 |
| Hemoglobin | 0.9122 | 0.7397 | 1.125 | 0.3906 |
| PB Platelet count | 0.9973 | 0.9921 | 1.003 | 0.3244 |

MDS, Myelodysplastic Syndrome; AML, Acute Myeloid Leukemia; VAF, Variant Allele Frequency; HR, Hazard Ratio

### Clinico-pathologic and genetic correlates of low and high EAp53 scores

Using the EAp53 score as a continuous variable, a higher EAp53 score positively correlated with a higher number of *TP53* mutations (p=0.00062), higher PB platelet counts (p=0.041), and higher serum fibrinogen levels (p=0.009). The EAp53 score negatively correlated with concurrent somatic mutations in *RUNX1* (p=0.038) and *EZH2* (p<0.001). There was no correlation between EAp53 score and PB/bone marrow (BM) blast percentage, *TP53* VAF or number of cytogenetic abnormalities.

Using the EAp53 as a categorical variable with a cut-off score of 52, low-risk patients had fewer cytogenetic abnormalities compared to those with high-risk EAp53 [median number 3 (range, 0-14) vs. 7 (range, 0-27); p=0.019]. High-risk EAp53 MDS had a significantly higher frequency of complex karyotype, defined here as 3 or more abnormalities (p=0.0241), and a monosomal karyotype (p=0.0043). There were no significant differences in the frequencies of normal karyotype, del(5q), del(7q) or del(17p) abnormalities. There were no significant differences in the distribution of IPSS-R risk categories, etiology (therapy-related) or treatment characteristics between the two groups.

In terms of *TP53* mutation characteristics, most *TP53* mutations involved the DNA-binding domain in both high- and low-risk EAp53 categories. High-risk EAp53 MDS patients had a higher number of concurrent *TP53* mutations compared to low-risk EAp53 patients (32% vs. 6%, p=0.027). High-risk EAp53 patients had a higher frequency of multi-allelic *TP53* alterations compared to low-risk EAp53 patients (63% vs. 29%, p=0.0087). There was no significant difference in the median VAF of *TP53* mutations between the two groups (34 vs. 23, p=0.68) (**Table 3**).

**Table 3.** Comparison of clinicopathologic characteristics between *TP53* mutated AML/MDS patients with low EA and high EAp53 scores. P-values were calculated using the two-sample t test (continuous variables), Fisher’s exact test (2 categories), or chi-squared test (more than 2 categories).

| Characteristics | Low EAp53 MDS | High EAp53 MDS | p-value |
| --- | --- | --- | --- |
|  | n=17 | n=253 |  |
| **Therapy-related** | 7 (41%) | 115 (45%) | 0.81 |
| **Age** | 72 (39-81) | 68 (18-90) | 0.92 |
| **Gender (M)** | 10 (59%) | 158 (62%) | 0.80 |
| **IPSS-R** |  |  | 0.28 |
| 1 | 0 (0%) | 9 (4%) | ns |
| 2 | 2 (12%) | 17 (7%) | ns |
| 3 | 3 (18%) | 16 (6%) | ns |
| 4 | 2 (12%) | 59 (23%) | ns |
| 5 | 10 (59%) | 152 (60%) | ns |
| **Karyotype** |  |  |  |
| Diploid* | 2 (12%) | 23 (9%) | 0.6628 |
| Complex* | 10 (59%) | 209 (83%) | 0.0241 |
| Non-complex | 7 (41%) | 44 (17%) | 0.0241 |
| del(5q)/ monosomy 5 | 6 (35%) | 148 (59%) | 0.4484 |
| del(7q)/ monosomy 7 | 5 (29%) | 116 (46%) | 0.4581 |
| Del(5q)/-5 & del(7q)/-7 | 4 (24%) | 84 (33%) | 1 |
| del(17p)* | 4 (24%) | 89 (35%) | 0.4333 |
| Monosomal karyotype | 10 (59%) | 222 (88%) | 0.0043 |
| **Number of cytogenetic abnormalities\*** |  |  |  |
| Median | 3 (0-14) | 7 (0-27) | **0.019** |
| **Number of *TP53* mutations\*** |  |  |  |
| 1 | 16 (94%) | 173 (68%) | **0.027** |
| 2 or more | 1 (6%) | 80 (32%) |  |
| ***TP53* allelic state** |  |  | **0.0087** |
| Monoallelic | 12 (71%) | 93 (37%) |  |
| Multi-allelic | 5 (29%) | 160 (63%) |  |
| **Mutation location\*** |  |  |  |
| DBD | 16 (94%) | 249 (98%) | 0.46 |
| OD | 1 (6%) | 2 (0.8%) |  |
| CTD | 0 | 1 (0.4%) |  |
| AT | 0 | 0 |  |
| TAD | 0 | 1 (0.4%) |  |
| **Median VAF for TP53 mutation** | 23 (1-94) | 34 (1-94) | 0.68 |
| **Treatment regimens** |  |  |  |
| Chemotherapy-based | 1 (6%) | 17 (7%) | 0.22 |
| Chemo + HMA-based | 2 (12%) | 6 (2%) |  |
| HMA alone | 6 (35%) | 105 (4%) |  |
| HMA + other | 2 (12%) | 55 (22%) |  |
| Others | 0 | 3 (1%) |  |
| None | 4 (24%) | 18 (7%) |  |
| Unknown | 2 (12%) | 49 (19%) |  |
| **Complete Blood Counts** |  |  |  |
| ANC | 1.0 (0.3-9.5) | 1.1 (0-19.8) | 0.89 |
| Platelet count | 42 (7-241) | 49 (2-481) | 0.86 |
| Hemoglobin | 9 (8-12) | 9 (6-16) | 0.64 |
| PB blast% | 4 (0-17) | 1 (0-25) | 0.31 |
| BM blast% | 6 (1-25) | 7 (0-28) | 0.51 |
| Dead | 7/17 (41%) | 185/253 (73%) | **0.010** |
| Transplant | 2/16 (12%) | 34/252 (13%) | 1 |
| OR | 7/9 (78%) | 98/171 (57%) | 0.31 |
| CR | 2/9 (22%) | 51/171 (30%) | 1 |

VAF, Variant Allele Frequency; IPSS, R-IPSS, revised International Prognostic Scoring System; HMA, hypomethylating agent; DBD, DNA-binding domain; TAD, Transactivation Domain; CTD; OD; AT

In order to assess the entire somatic mutation spectrum, we compared mutation characteristics among patients that underwent sequencing using a uniform next-generation sequencing (NGS) gene panel. The median number of somatic mutations in low- and high-risk EAp53 patients (including *TP53*) was 3 and 1, respectively (p=0.000002). A higher proportion of low-risk EAp53 patients had mutations in additional genes (63% vs. 33%; p=0.05) compared to those with high-risk EAp53 (**Figure 2A**), and these involved *NRAS* (p=0.02) and *RUNX1* (p=0.02) genes. Low-risk EAp53 patients also tended to have a higher frequency of mutations in *NPM1*, *WT1*, and *ASXL1* genes compared to those with high-risk EAp53 (**Figure 2B**). There was no difference in molecular composition between MDS and oligoblastic AML subgroups.

**Figure 2.**
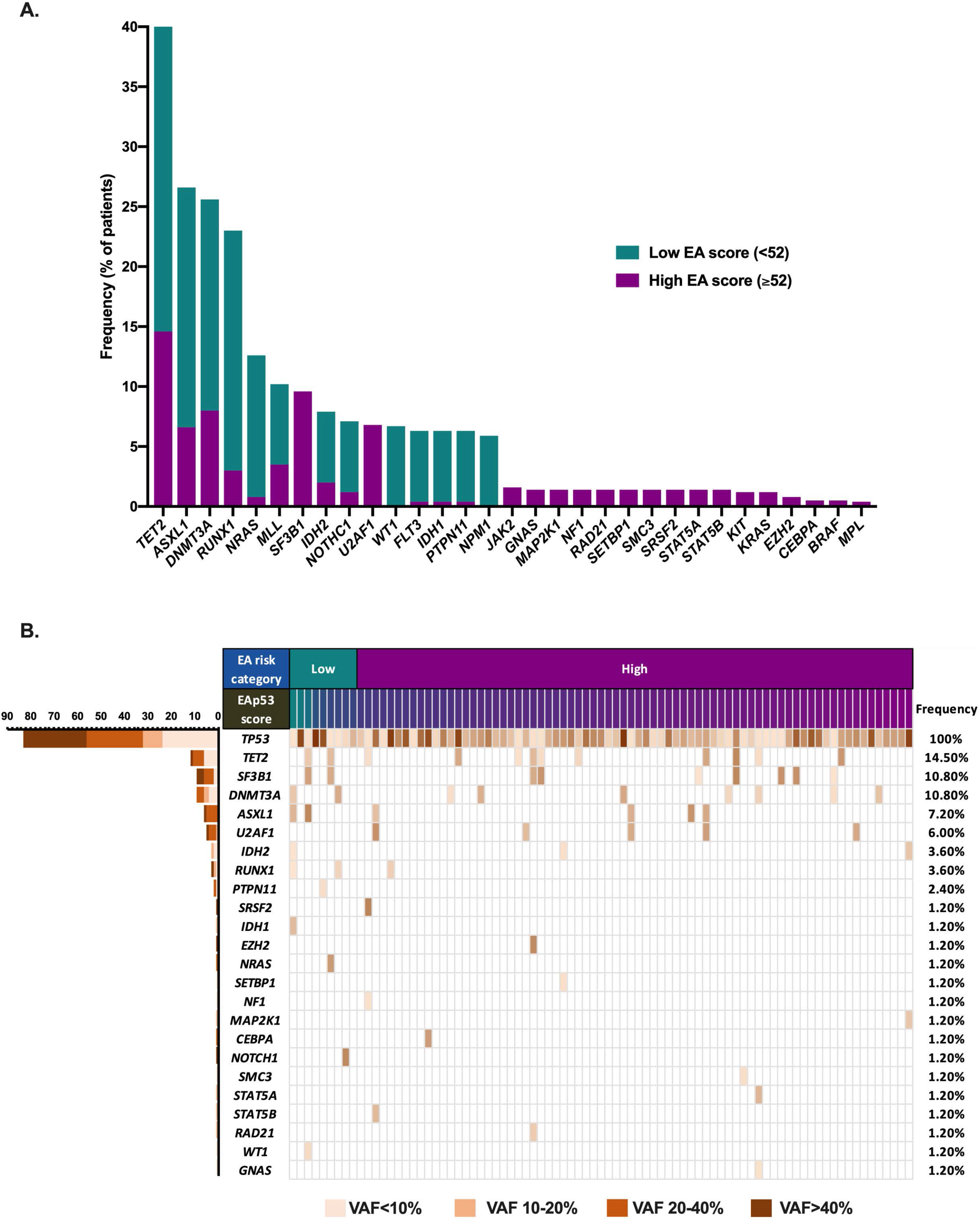
**(A)** Mutational frequencies of genes in the cohort separated by EAp53 risk category. Low-risk EAp53 MDS patients had a significantly higher frequency of mutations in *NRAS* and *RUNX1*. **(B)** The mutation spectra of low and high EAp53 AML/MDS patients were different; high EAp53 patients lacked additional mutations in other genes (median of 1 mutation per case) while low EAp53 patients had a median of 3 additional mutations. High EAp53 patients had more frequent chromosomal aberrations.

Unlike wild-type TP53 protein that undergoes rapid MDM2-mediated degradation, mutant TP53 protein accumulates in tumor cells that can be detected by immunohistochemistry (IHC). To assess the possible downstream effect of the EAp53 score of the *TP53* mutation on the translated protein, we performed TP53 IHC on BM biopsy specimens (n=30). For this, we included only patients with a single missense *TP53* mutation. We quantified the IHC expression using an H-score (based on a combination of percent positive cells and intensity of expression), which showed a significant correlation with EAp53. The median H-score for TP53 protein expression by IHC was significantly different between wild-type (median 6.4, n=4), low-risk EAp53 (median H-score 47.5; median EA score: 27.9; median VAF: 30; n=10) and high-risk EAp53 (median H-score 157.5; median EA 84.8, median VAF: 48; n=20) [wild-type vs. low EA, p=0.04; low vs. high EA, p=0.0014] (**Figure 3A**). H-score also correlated with *TP53* VAF (p=0.00015; rho (ρ)=0.61). These findings provide support that EAp53 score reflects functional consequences of clinical importance in patients with AML/MDS. Representative images of TP53 IHC staining are shown in low-risk (**Figure 3B, 3C**) and high-risk (**Figure 3D**) EAp53 cases.

**Figure 3.**
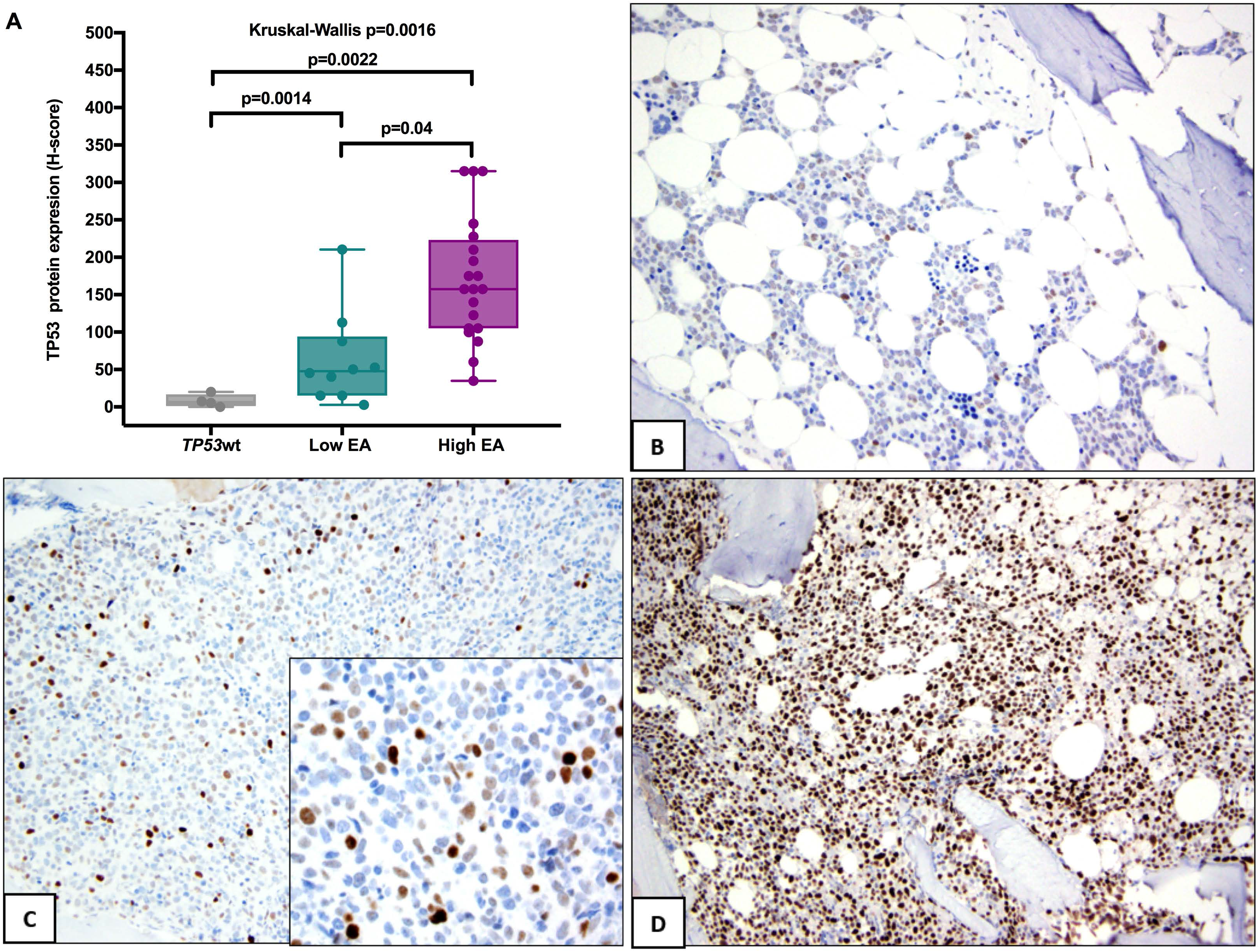
**(A)** Representative images showing downstream assessment of TP53 protein expression by immunohistochemistry (IHC). *TP53* wild-type, *TP53* mutant low and high EAp53 groups showed significant differences in the median H-scores for TP53 protein expression by IHC. Immunohistochemical staining patterns of low-risk and high-risk EAp53 MDS patients are shown in **B-D**. IHC staining patterns in MDS with low-risk EAp53 correlated with a lower H-score: **(B)** diploid karyotype (weak staining in ~80% of cells) and **(C)** complex karyotype (dual population of cells: strong positive cells in ~5% and weak positive cells in 10% of all cells; inset shows staining at 1000x magnification). **(D)** IHC staining pattern in MDS with high EAp53 mutation showing a high H-score (strong positivity in >50% of cells).

### Mutation mapping on TP53 protein structure sites

To correlate survival with the molecular and structural impact of the TP53 mutations in this patient cohort, we used a crystal structure of the TP53 core domain in complex with DNA (PDB ID: 4HJE; see Methods). Per the Evolutionary Trace analysis, most of the important TP53 residues were located near the DNA binding site or within the protein structural core and solvent inaccessible, as shown in **Figure 4A**. The *TP53* mutations of this patient cohort mapped to the evolutionarily important sites of the TP53 core domain (**Figure 4B**). However, for certain residues, despite the similar Evolutionary Trace importance, patients had variable survival outcome (**Figure 4C**). To determine whether this difference could be attributed to the location of the mutants in the TP53 structure, we divided the patients into two groups based on a survival cut-off of 10 months and accordingly annotated each *TP53* mutation. The TP53 variants associated with poor survival (OS <10 months) in patients formed two clusters: a large cluster interfacing with the DNA binding site and a small cluster formed by residues V157, Y220, L257 and E258 (**Figure 4D**). Using more extreme cut-offs (OS <5 months and >15 months, OS <2.5 months and >20 months, OS <1 month and >30 months) yielded the same results. These data suggest that structure location can inform us regarding the expected survival of MDS patients and may help to further stratify the high-risk group defined by the EAp53 score.

**Figure 4.**
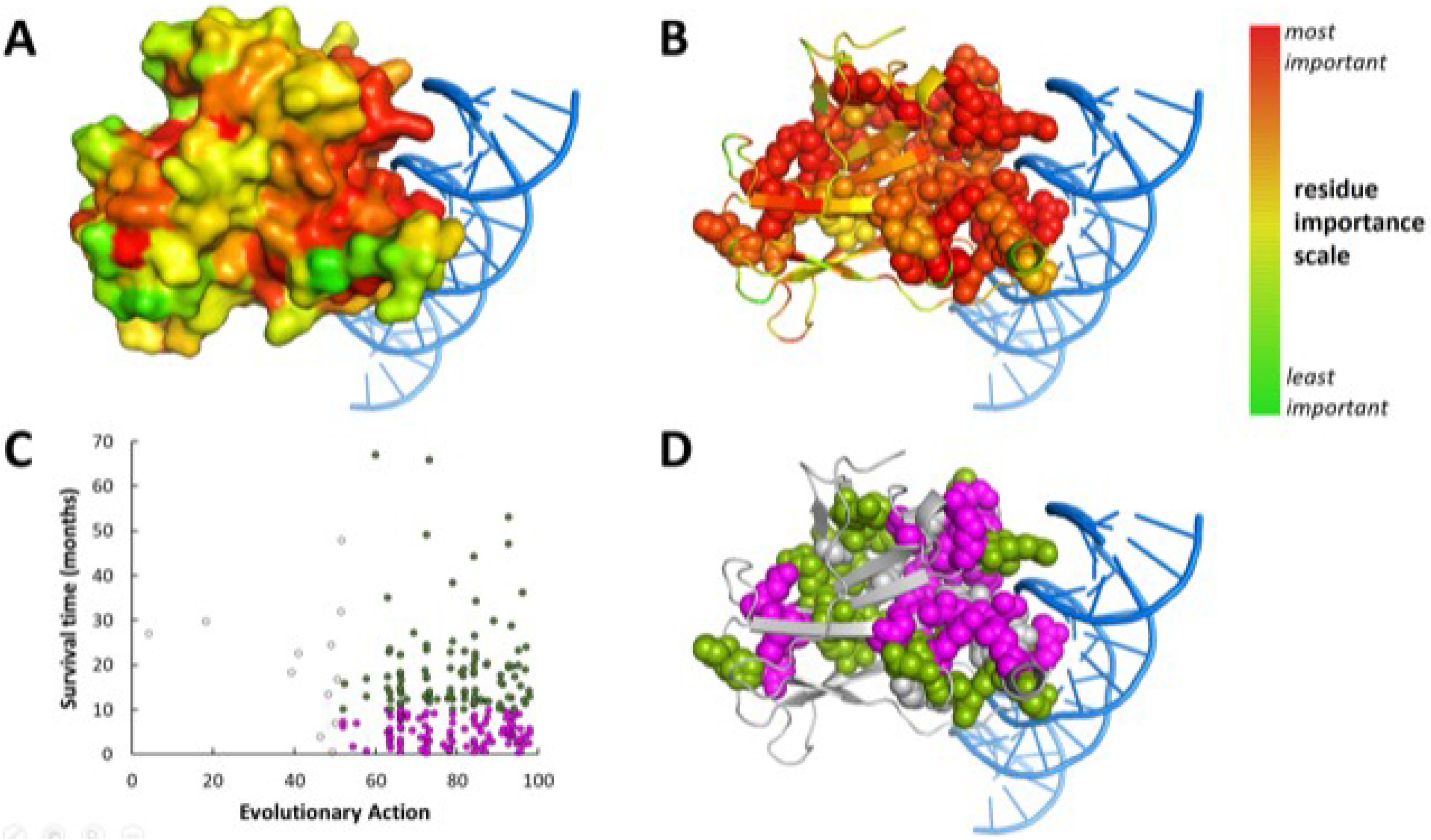
Analysis of TP53 protein structure, function, and patient survival, using the TP53 core domain structure bound to DNA (PDB ID of 4HJE, visualized by PyMOL). **(A)** A surface representation of the TP53 structure, colored according to the importance of the TP53 residues as estimated by the Evolutionary Trace (ET) method (red represents the most important residues and green the least important ones). **(B)** A cartoon representation of the TP53 structure, colored again according to the ET importance, with the residues that have at least one mutation in our patient cohort shown with atomic spheres. **(C)** A survival time versus the Evolutionary Action plot, for 215 patients who were divided into 113 patients with poor survival (pink color) and 102 patients with good survival (green color) using a threshold of 10 months (see Methods). **(D)** A cartoon representation of the TP53 structure with residues mutated mostly in patients with poor survival represented by pink atomic spheres, residues mutated mostly in patients with good survival represented by green atomic spheres, and residues with equal numbers of patients with poor and good survival represented by white atomic spheres.

### Mutational dynamics during disease evolution and therapy

We assessed the stability of missense *TP53* mutations with low and high EAp53 scores by serial sequencing using an NGS assay that covers the whole coding region at 1% detection limit. Nine patients with low EAp53 and 36 patients with high EAp53 scores underwent repeat NGS sequencing at one additional time point.

All but one low-risk EAp53 patients had 1 *TP53* mutation at baseline. Among them, partial response (PR) or hematologic improvement (HI) was noted in 3 of 9 (33%) patients [none achieved complete remission (CR/CRi/mCR], 2 (67%) of which showed clearance of *TP53* mutations (**Figure 5A**). Five patients either did not respond (**Figure 5B**) or transformed to AML (**Figure 5C and D**), including one that relapsed following allogeneic stem cell transplantation (**Figure 5E**), and 4 showed persistence of the *TP53* mutant clone. In 1 patient, a low-risk EAp53 *TP53* mutation (p.N239D) was detected in the PB at the time of diagnosis of a solid tumor with a different *TP53* mutation (p.C141Y) (clonal hematopoiesis). Eight months following chemotherapy, this patient progressed to therapy-related MDS with *TP53* p.N239D and other gene mutations. Subsequently, while undergoing therapy with decitabine, the *TP53* mutation disappeared with acquisition of *KRAS* and additional *RUNX1* mutations (**Figure 5F**). Four out of 5 (80%) patients who progressed to AML acquired additional mutations in *NRAS, KRAS/RUNX1, IDH1* and *JAK2*. No additional gene mutations were acquired at response. None of the low-risk EAp53 patients acquired additional *TP53* mutations.

**Figure 5.**
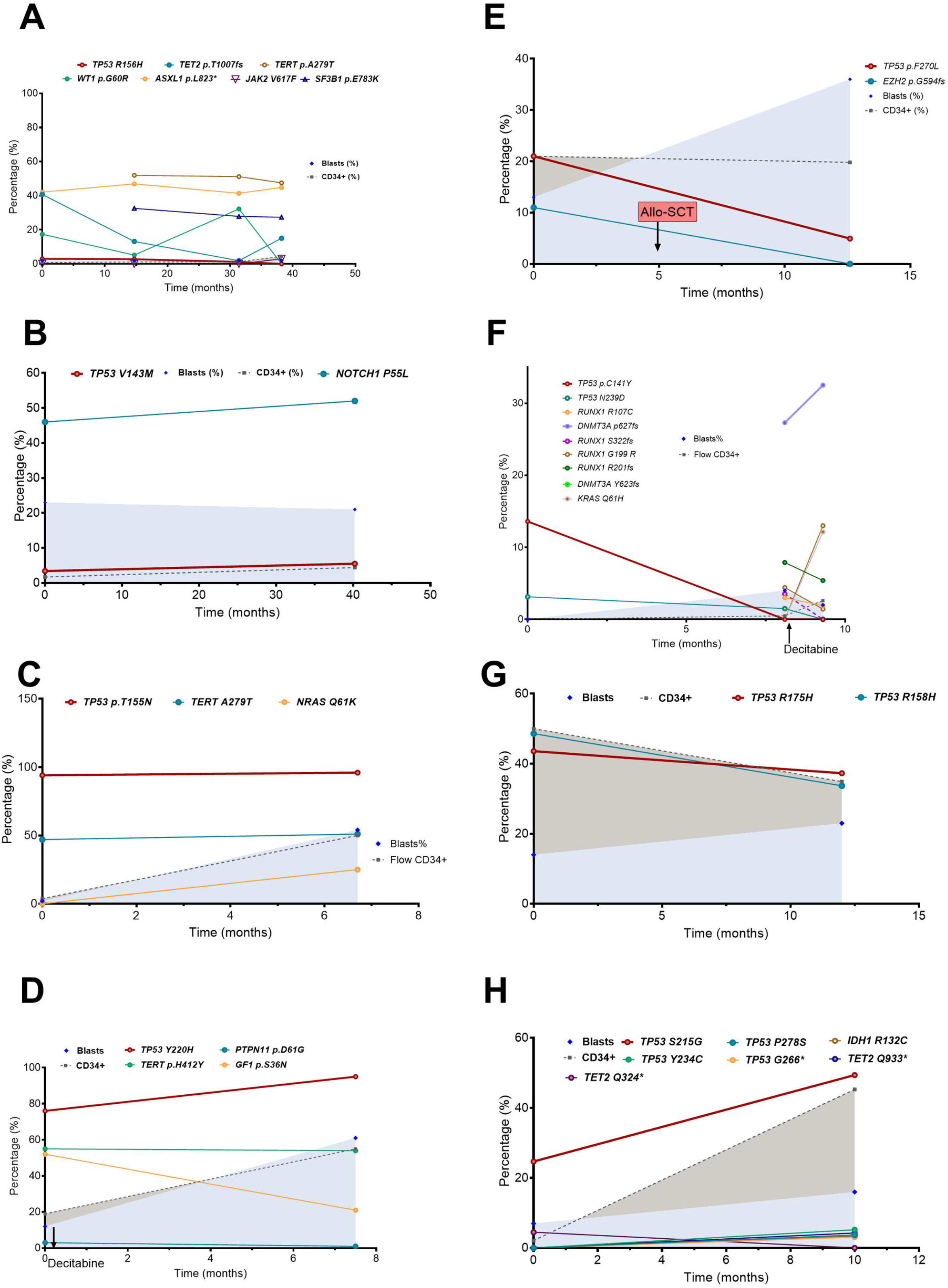
Sequential analysis of low-risk versus high-risk EAp53 *TP53* mutated patients showed higher mutations clearance rates in low-risk EAp53 versus high-risk EAp53 MDS patients. **(A)** Clearance of low-EA risk *TP53* mutation at response (HI). **(B)** Persistence of low-risk EA *TP53* mutation in MDS with no response. Persistence of low-risk EAp53 *TP53* mutations with additional co-mutations at AML transformation with **(C)** and without **(D)** increase in *TP53* VAF. **(E)** Recurrence of the same low-risk *TP53* mutation post-allogeneic stem cell transplant at the time of relapse. **(F)** Clearance of low-risk *TP53* mutation identified at the time of clonal hematopoiesis that evolved to therapy-related MDS with decitabine therapy. **(G)** Persistence of high-risk EA *TP53* mutations in a patient that did not respond to HMA treatment. **(H)** An MDS patient with high-risk EAp53 *TP53* mutation showing clonal evolution with additional *TP53* mutations at the time of AML transformation.

Patients with high-risk EAp53 *TP53* mutations included: 29 patients with 1 mutation and 7 patients with 2 mutations at baseline. 6 (17%) patients transformed to AML. Among 11 (27%) patients who achieved CR/CRi/mCR, the *TP53* mutation disappeared in 5 (45%) patients. In the remaining 25 (70%) patients who did not achieve CR/CRi/mCR, the original *TP53* mutation persisted (**Figure 5G**). One patient acquired 3 additional *TP53* mutations at the time of AML transformation [2 point mutations, both with high EAp53 scores, and 1 nonsense mutation] (**Figure 5H**). Only 2 (8%) patients acquired additional mutations at the time of AML progression, 2 with *NRAS* mutations, 1 with an *IDH1* mutation and 1 with concurrent *IDH1* and *TET2* mutations. No additional mutations were observed at response.

These findings led us to conclude that the specific *TP53* variant remains stable throughout the disease course. Additional *TP53* mutations acquired by a small proportion of high-risk EAp53 patients also had a high-risk EAp53 score. In contrast, low-risk EAp53 MDS patients often acquired mutations in other genes at the time of progression or AML transformation. The only exception was one patient: t-MDS patient (with a history of DLBCL treated with chemotherapy) with R248Q (high-risk EAp53) at <10% VAF at diagnosis was treated with azacitidine. Following allogeneic SCT, the patient developed an N235S mutation (low-risk EAp53: 39.26) at 10 months follow-up. Based on the absence of morphologic disease, 100% engraftment by chimerism assay, this mutation was likely of donor origin.

## DISCUSSION

The distinct functional consequences of various *TP53* mutations impact patient outcomes and responses to therapy. Here we utilized the evolutional action scoring system (EAp53) to quantify the deleterious effects of missense *TP53* mutations. Within a large, well-characterized cohort of 270 patients with newly diagnosed MDS and oligoblastic AML with missense *TP53* mutations, we demonstrated that EAp53 scores (<52) identified a small subset of patients with a favorable outcomes. In addition, clustering of high-risk EAp53 mutations with worse survival within TP53 protein structure suggests that the location of the mutations can further stratify high-risk EAp53 mutations.

EAp53 is a computational score to measure the deleterious impact of missense *TP53* mutations. EAp53 score is a single score based on 2 components. The first component is based on the functional sensitivity of the sequence position (example, codon 248) to sequence variation. This is calculated using the extensively validated “evolutionary trace” method to identify key functional/ structural protein residues. Here, each sequence position is assigned an ET grade based on the degree of phylogenetic divergence (so, larger ET grade, means more “evolutionary sensitive” to sequence variation due to larger phylogenetic divergence). The second component of the EAp53 score is the perturbation size of the specific amino acid substitution (example, arginine to tryptophan switch in p.R248W). So, the mutational impact should be greater when the mutated residues are more evolutionarily sensitive to sequence variations and also when the amino acid change is least conservative. The two components are computed into a single EAp53 score following normalization that represents the percentile rank of each variant within the protein. So, EAp53 score of 84.11 (for p.R248W) means, that the phenotypic impact of this variant is larger than 84.11% of random AA changes in the protein. EAp53 score ranges from 0-100, with higher scores indicating a higher impact on TP53 transcriptional activity and protein function and 0 indicating wild-type TP53 function. ^32, 33^.

To the best of our knowledge, none of the low-risk EAp53 *TP53* mutations described here were known polymorphisms. All mutations were carefully curated based on evidence from literature and data from our institutional database. None of the variants were identified in the SNP (single nucleotide polymorphism) databases such as dbSNP or EXAC, and were annotated in COSMIC as somatic. Nine of 17 cases had repeat NGS sequencing, which showed significant variations in the *TP53* mutant VAFs or clearance in contrast to polymorphisms, strongly suggesting the somatic nature of the mutations.

Our findings show that *TP53* EAp53 score is independently prognostic in MDS. So far, studies have attributed the heterogeneity in outcome to mutation characteristics such as VAFs and associated parameters such as the presence of CK and concurrent 17p deletion ^5, 7, 9, 10, 11, 12, 13, 14, 15^. In this cohort, high EAp53 score correlated with CK/MK but not with *TP53* VAF. By multivariante analysis, EAp53 risk was independently prognostic for OS in addition to the IPSS-R scores. Notably, although associated with worse OS, *TP53* VAF did not show independent prognostic value. It is important to note that the study group is unique since it only includes missense *TP53* mutations and excludes nonsense and frameshift mutations; the latter are likely to have a higher VAF due to a loss-of-function phenotype. The findings with respect to VAF and outcome in MDS have not been consistent. A few studies reported that high *TP53* VAF (>40%) predicted for complex cytogenetics and worse OS ^5, 7, 14^ but others did not ^9^. Of note, the VAFs in the current study have not been normalized based on copy number change.

To translate this finding into clinical practice, both VAF and karyotype yield are affected by the adequacy of the aspirate material and the degree of hemodilution. *TP53* mutated MDS is frequently associated with fibrosis and dry tap ^37^, and the yield may not always be sufficient for analysis. Here we show that EAp53 score is a biologically stable parameter. In our study, *TP53* mutation(s) at baseline were stable throughout the disease course; a majority of the patients did not acquire any additional *TP53* mutations. Given that VAF can dynamically change through the course of therapy, we suggest that EAp53 score may be a more stable predictive biomarker of outcome for baseline assessment and risk-stratification. In our study, the median VAF of *TP53* mutation was not different between the 2 groups with distinct prognoses.

We determined an EAp53 cut-off score based on outcome, similar to studies conducted in head and neck cancers. We identified distinct clinical, cytogenetic, and mutation characteristics in the 2 different categories, thereby supporting the clinical significance of this scoring system. In addition, several important conclusions can be inferred from these findings. First, high-risk EAp53 was associated with CK, MK and higher number of *TP53* mutations. This suggests that the type of *TP53* mutation is the primary driver and dictates the degree of karyotypic and genomic complexity. Second, in contrast to high EAp53 MDS, low EAp53 MDS and AML patients had at least 1 additional concurrent gene mutation (frequently in *NRAS*), supporting the role of RAS pathway activation in leukemogenesis. The need for additional hits in high-risk EAp53 patients may be abrogated by the higher frequency of chromosomal aneuploidies, frequently noted in chromosomes 17 and 5, that harbor negative regulators of the RAS pathway ^38^. Thus, low EAp53 *TP53* mutations may not be the sole drivers of leukemia pathogenesis, and these additional hits could potentially modify the phenotype and outcome. It is unlikely that low and high EAp53 subsets reflect different positions on the trajectory from early to late disease. The VAFs of mutations (*TP53, NRAS* and *RUNX1*) is low-EAp53 patients were high at onset, and the additional somatic mutations did not disappear with time. None of the high-EAp53 patients developed additional mutations later in the disease course.

In this study, we confirmed the differential downstream effects of low and high EAp53 mutations in clinical BM samples using IHC to assess protein expression. Since percentage positivity depends on VAF, we used an H-score (percentage positivity multiplied by intensity) to evaluate protein expression. Patients with a high EAp53 had a higher H-score compared to those with low EAp53 scores. These results are in accord with the mRNA expression profiling studies in head and neck squamous cell carcinoma cells, where high-risk EAp53 mutations had marked reduction in wild-type TP53 function while low-risk EAp53 cells had residual TP53 function ^25, 26^. TP53 IHC is a rapid and reliable test for detecting the presence of *TP53* mutations in BM specimens for routine clinical care with a high degree of sensitivity and specificity. The degree of TP53 IHC positivity is prognostic in myeloid malignancies ^39, 40, 41^. In addition, IHC can inform the EAp53 risk of the *TP53* mutation.

The findings in this study are important in lieu of novel therapeutic strategies to overcome *TP53* mutated malignancies. *TP53* mutated clones are inherently resistant to genotoxic stress. Treatment with conventional chemotherapy leads to poor response and a high likelihood of relapse by clonal selection ^42, 43, 44, 45^. *TP53* mutant clones in MDS and AML patients are particularly sensitive to hypomethylating agents, however, responses are short-lived ^46^. The diversity in *TP53* mutations with distinct functional effects necessitates *TP53* mutation type-specific therapeutic strategies ^19, 47^. In low-risk EAp53 mutants, strategies to utilize the residual TP53 function might be beneficial while approaches such as small molecules that target missense *TP53* mutations to restore wild-type function as a result of protein structure modification may be appropriate for high-risk EAp53 mutants. It is possible that the efficacy of these small molecules may depend on the location of the mutations on the TP53 protein. These are currently under investigation and include: (a) APR-246 to restore mutant TP53 function, which has shown efficacy in phase I/IIa clinical trials in leukemia patients and has recently received break-through designation by the FDA in MDS ^28, 29^(b) COTI-2, with activity against both mutant and wild-type TP53 [1]. Thus, EAp53 score and structural mapping could potentially be used to select and guide personalized treatment to improve clinical outcomes.

Our study has limitations. This is a retrospective study of patients with missense *TP53* mutations at the time of diagnosis. Our study did not evaluate truncating variants, nonsense mutations and splice-site variants. We did not use either chromosomal microarray or capture-based NGS techniques to accurately determine *TP53* copy number changes. Despite being one of the largest *TP53* mutated study cohorts, the relatively low frequency of low EAp53 MDS (~6%) warrants validation of the cut-off criteria in larger multi-center cohorts. Such cohorts may also be used to validate our evidence that protein structural considerations can help in further stratification of the high-risk EAp53 group.

In summary, this study shows the clinical utility of EAp53 score in a well-characterized cohort of *TP53* mutated MDS patients treated primarily with hypomethylating agents. EAp53 score, a quantitative measure of the degree of TP53 functional defect, associates with specific clinico-pathologic characteristics. A subset of *TP53* mutated MDS patients with low (<52) EAp53 score associated with improved outcomes independent of VAF or IPSS-R risk scores and paucity of complex multi-allelic *TP53* alterations. EAp53 is a stable biological parameter that is not influenced by either therapy or time, and can be used for baseline risk assessment and treatment decisions.

## METHODS

### Patient selection

All patients presenting to The University of Texas MD Anderson Cancer Center between January 1, 2012 and December 31, 2018 with a new diagnosis of MDS or AML with <30% blasts, defined as MDS with excess blasts in transformation per the National Cancer Center Network (NCCN) guidelines^48^, with at least 1 missense mutation in *TP53* were included in the study. The reasons for the inclusion of oligoblastic AML (20-29%) were as follows. First, the NCCN recognizes this as “MDS-EB in transformation” to be within the spectrum of MDS based on similarity in clinical course^48, 49^. The currently validated prognostic models for MDS including IPSS-R incorporate this subgroup of patients^50^. Second, the current understanding of genomic data suggests that mutated MDS/oligoblastic AML represent the spectrum of the same disease, and a cut-off of 20% seems arbitrary^51^. Hence, we expanded the blast cut-off to 30% for the purpose of this study similar to the International Working Group for Prognosis in MDS (IWG-PM). All patients underwent morphologic evaluation of BM using World Health Organization criteria^51^. Conventional G-banded karyotyping analysis was performed from 24-hour and 72-hour cultures of bone marrow aspirate using standard techniques and reported using International System for Human Cytogenetic Nomenclature 2013^52^. The prognostic risk was calculated using R-IPSS^50, 53^. Only patients with at least 1 missense *TP53* mutation at the time of diagnosis (not those that developed over the course of the disease) were included. Informed consent was obtained and all studies were performed based on the institutional approved protocols in accordance with the Declaration of Helsinki.

### Molecular analysis

All patients underwent *TP53* mutation analysis by NGS based multi-gene panels within the Clinical Laboratory Improvement Amendments (CLIA)-certified Molecular Diagnostics Laboratory, as previously described^54^. A majority of patient samples were analyzed with either 28-gene or 81-gene panels that included the entire coding region of the *TP53* gene. Briefly, the genomic DNA extracted from whole mononuclear cells from fresh BM aspirates underwent amplicon-based targeted NGS analysis on a MiSeq sequencer (Illumina Inc., San Diego, CA, USA), as described previously^52, 55^. GRCh37/hg19 was used as a reference for sequence alignment. A minimum of 1% VAF with adequate coverage was required for variant calling. Since matched germline samples were not sequenced, the somatic nature of the variants was inferred based on the VAFs, evidence from the literature and online databases such as COSMIC and data from our institutional cohort. Variants reported in the Exome Aggregation Consortium [ExAC], dbSNP 137/138, and 1000 Genomes Project databases were excluded^56^. *FLT3* ITD mutations were evaluated by PCR-based capillary electrophoresis.

### EAp53 scoring

EAp53 scores for all missense *TP53* mutations were obtained at the EAp53 server at http://mammoth.bcm.tmc.edu/EAp53 using the Evolutionary Trace approach. Higher scores represent alterations that are more deleterious. For patients with more than 1 *TP53* missense mutations, the highest EAp53 score was considered irrespective of the VAF. The optimal cut-off EAp53 score for distinguishing a low versus high EAp53 score for *TP53* mutations in MDS was obtained as described in Results.

### Immunohistochemical staining for TP53 protein expression

TP53 IHC was performed using monoclonal anti-TP53 antibody clone DO-7 (Dako, Carpinteria, CA, USA) using standard techniques described elsewhere^57^ on whole BM biopsy sections in a CLIA-certified laboratory. TP53 IHC staining was scored independently by two hematopathologists (RK-S, CB-R) in a blinded fashion. In addition to percent cells showing nuclear positivity of all nucleated cells (a total of at least 5000 nucleated cells were counted), cases were also semi-quantitatively scored for intensity (negative=0/1; weak=2; moderate=3; strong=4). An H-score was calculated by multiplying percent positivity and the intensity scores. In cases showing dual populations of cells with different intensities, an average H-score from both populations was calculated.

### Statistical analysis

Responses were evaluated using the 2006 International Working Group criteria^58^. OS was defined as the number of months from the time of diagnosis to death or last follow-up. The time to transformation was calculated for WHO-defined MDS patients as the number of months from diagnosis to AML. Relapse-free survival (RFS) was defined as the number of months from the time of response until relapse or death. Transformation-free survival (TFS) was defined as the number of months from diagnosis to AML transformation or death. Patients who were alive at their last follow-up were censored on that date. The median (OS, RFS and time to AML transformation were evaluated by the Kaplan-Meier method. Univariate Cox proportional hazards analyses were used to identify associations between risk factors and survival. In addition to clinico-pathologic parameters, *TP53* VAF and EAp53 scores were correlated to clinical outcomes. The optimal EAp53 cutoff was determined and through the use of recursive partitioning and regression trees (RPART, based on the CART algorithm)^59, 60, 61^. Multivariate Cox proportional hazards analyses were also conducted within each diagnosis, including additional clinico-pathologic risk factors and *TP53* VAF. Logistic regression (for continuous variables) and Fisher’s exact test (for categorical variables) were used to study the association of overall response rates (ORR) or complete response rates (CRR) and risk factors. Statistical analysis was performed using R version 3.5.1^62^.

### 3D Structure analysis of the TP53 protein

The structural analysis of the TP53 protein was conducted using the PyMOL molecular visualization system and the crystal structure of the TP53 core domain in complex with DNA (PDB ID of 4HJE)^63^. The importance of the protein residues was estimated by the Evolutionary Trace approach^31^ and was represented on the structure using a color scale, red (most important) to green (least important), using the PyETV plugin^64^. Next, to correlate survival with molecular structural location, we performed structural analysis on patients with high-risk EAp53 (>52) mutations. After excluding patients who were alive with <10 months follow-up time (n=43), we divided 215 high-risk EAp53 score patients into 2 groups based on survival cut-off of 10 months: 113 with a survival of <10 months and 102 patients with a survival of ≥10 months. Based on this data, each TP53 protein residue was considered as high- or low-risk when >50% of the patients had poor or good survival, respectively. Residues with equal numbers of patients with poor and good survival were considered “neutral”. Using more extreme cut-offs (<5 months and >15 months, <2.5 months and >20 months, <1 month and >30 months) yielded the same results.

---

**Supplemental Table 1.** Comparison of survival outcomes of *TP53* mutated MDS patients with low and high risk EAp53 scores

|  | **Low EAp53**<br>Median (95% CI) | **High EAp53**<br>Median (95% CI) | **p-value** |
| --- | --- | --- | --- |
| OS* | 47.8 (13.3-NA) | 10.0 (8.8-12.2) | **0.013** |
| RFS* | 12.6 (4.0-NA) | 6.6 (5.3-8.4) | 0.20 |
| TFS* | 93.8 (NA-NA) | 49.5 (44.8-60.6) | **0.05** |
| ORR** | 7/9 (78%) | 98/171 (57%) | 0.31 |
| CRR** | 2/9 (22%) | 51/171 (30%) | 1 |

\*p-value calculated by log-rank
\*\*p-value calculated by Fisher's exact test
CI, confidence interval; MDS, Myelodysplastic Syndrome; OS; RFS; ORR; CRR

